# Glyphosate resistance evolution to lethal and sublethal doses in chemostat populations of model organism *Chlamydomonas reinhardtii*

**DOI:** 10.1101/2024.02.26.582134

**Authors:** Erika M. Hansson, Dylan Z. Childs, Andrew P. Beckerman

## Abstract

Herbicide resistant weeds are an increasing economic and ecological problem world-wide. Evolutionary theory and insight from experiments testing this theory are now a central part of solving resistance problems. More specifically, experimental evolution, where populations are allowed to evolve under specific conditions, can offer substantial insights into the trade-offs that govern the pace at which resistance arises. Here we leverage the green alga *Chlamydomonas reinhardtii* facing glyphosate as a model plant system to evaluate such theory, monitoring the level of evolved resistance and associated fitness costs throughout the course of adaptation. On a gradient of lethal and sub-lethal doses of glyphosate, we found evidence for evolved resistance from standing genetic variation but limited evidence for classic growth rate trade-offs that are expected to affect the pace of resistance evolution.

## Introduction

The widespread and persistent use of herbicides worldwide has resulted in a strong selective pressure for resistant phenotypes[1]. To date, 269 different weed species have evolved resistance to 167 different herbicides[2], all while the rate of discovery of new herbicide modes of action (functional changes at a cellular level in response to exposure) is declining[2, 3]. This is a growing problem, both ecologically and economically. Weeds are already responsible for ca. 10% of crop yield loss worldwide and with evolved resistance to herbicides constitute a higher potential threat to crop yield than any other pest[4].

Resistance is an evolutionary process, and understanding the evolutionary ecology of herbicide resistant weeds at both an organism and population level is pivotal to enable sustainable use of herbicides[5, 6]. The evolutionary dynamics of herbicide resistance have thus emerged as a central part of the toolset for managing it, centering on the fitness consequences of resistance in the presence and absence of herbicides[7, 8]. Resistance is predicted to confer theoretical intrinsic fitness costs based on trade-offs with normal cell function or resource investment that decreases resources available for growth or reproduction[1, 9]. As a population evolves resistance to herbicide its performance while herbicide-exposed increases as well as the minimum inhibitory concentration (MIC) of the herbicide, resulting in a flatter dose-response slope (Fig 1). An associated change in performance in the ancestral environment would thus be evidence of an intrinsic fitness cost. Resistance management strategies are largely based on the expectation that resistance will incur a sufficient fitness cost that the absence of herbicide will put it under negative directional selection[8]. However, costs are rarely found to be universal and often have an extrinsic, ecological component through compromising performance in specific environments[10] meaning their effect on the pace of evolution may be hard to estimate without empirical evidence to underpin the theory. Furthermore, the dynamics of the adaptation process will depend on the strength of the selective pressure, i.e. the herbicide dose applied[11, 1], but as the majority of studies focus on weed populations experiencing high doses designed to kill the vast majority of the population, the effects of lower doses are poorly understood[12, 13, 14]. Experimental evolution under tightly controlled conditions in the lab allows us to connect when shifts in resistance level and performance emerge with the overall pace and dynamics of evolution by monitoring resistance evolution in action with continuous testing of both the level of resistance and intrinsic fitness costs in the ancestral environment.

**Figure 1:**
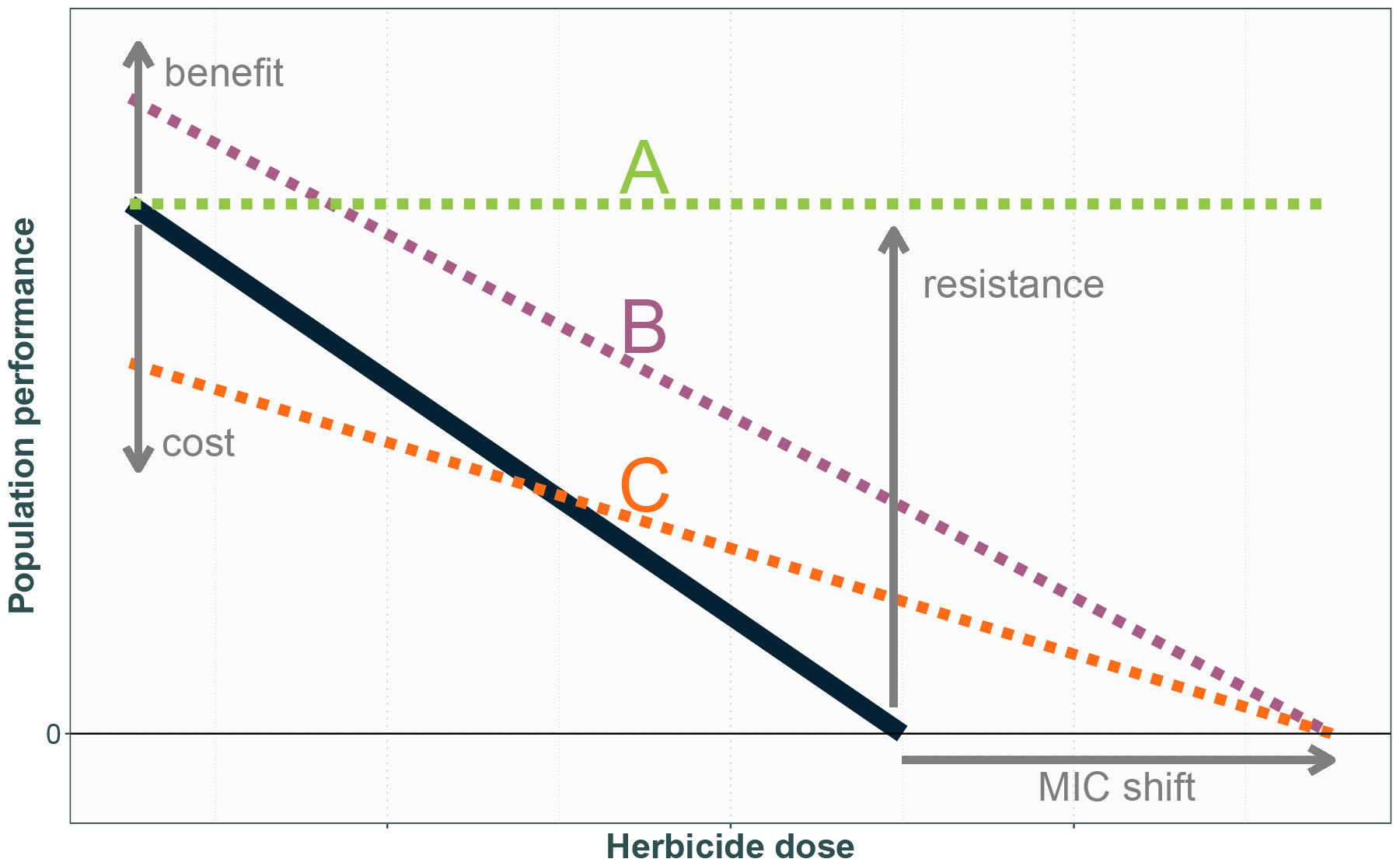
As the ancestral population represented by the black line evolves resistance to herbicide its performance while herbicide-exposed increases as well as the MIC of the herbicide, resulting in a flatter dose response slope, possibly associated with a change in performance in the ancestral environment. Examples of evolutionary outcomes are represented by the dotted lines: A has evolved resistance to herbicide at no cost to performance in the ancestral environment, and the magnitude of increased performance at high doses suggests a large shift in the MIC; B has evolved increased resistance to herbicide but with a slope suggesting a smaller shift in the MIC with a fitness benefit to performance in the ancestral environment; C has evolved increased resistance but this confers a cost in the ancestral environment.

Glyphosate has since the introduction of glyphosate resistant crops become the world’s most commonly used herbicide, and is thus of particular economic importance[15]. Due to the widespread use of glyphosate, documented resistant strains of weeds have rapidly increased in number[2] and become a serious problem for agriculture, as well as a major source of pollution for non-target ecosystems[16]. Glyphosate blocks the shikimate pathway by competitively binding to enzyme 5-enolpyruvylshikimate-3-phosphate synthate (EPSPS), outcompeting intended substrate phosphoenolpyruvate (PEP) and hindering aromatic amino acid synthesis[17]. Secondary effects on other cellular processes have also been documented, including disruption of photosynthesis and increased production of reactive oxygen species[18, 19, 20]. However, while the glyphosate mode of action is relatively well understood, we have little insight into how fundamental evolutionary theory regarding fitness costs drive pace and direction of evolving glyphosate resistance; the level of resistance and the costs of resistance appear largely dependent on the species and the specific molecular mechanism[9, 21]. Furthermore, factors like trait dominance[22], genetic background[23] and life history stage[24] as well as other stressors present in the environment like temperature[25] or competition[26] have a considerable effect on the phenotype. In some cases, resistance appears to be fitness neutral[27] or even confer a fitness benefit[28].

Here we experimentally evolve resistance to a lethal and a sublethal dose of glyphosate in populations of unicellular green alga *Chlamydomonas reinhardtii* using continuous flow-through chemostats. We use this system to evaluate the theory of intrinsic costs underpinning the pace of evolution of resistance. Using chemostats rather than batch cultures allows the populations to be kept at a steady state of constant exponential growth, removing effects of incidental nutritional stress or over-dilution as well as evolutionary bottlenecks. With its short generation time, small size and sensitivity to multiple commonly used herbicides[29, 30], *C. reinhardtii* has already been used to study efficacy of resistance management strategies[31, 32], characterising fitness costs[28], comparisons of evolutionary potential in communities of algae[33] and molecular analysis of herbicide resistance mutations[34, 35]. Furthermore, algal populations worldwide are still affected when herbicide contamination reaches non-target ecosystems through agricultural run-off[16] and as key primary producers their ability to evolve resistance and associated fitness costs may have far reaching consequences[36, 37]. While the ability for many algal species to evolve resistance to herbicide pollution is well documented, understanding the dynamics of that process with any associated trade-offs in fitness is an important missing piece of the puzzle to be able to understand the effects pollution may have on ecosystems and community assemblages[33, 38].

We hypothesise that both glyphosate treatments should cause growth-inhibition leading to continuous population decline and that the evolution of resistance should thus lead to population density resurgence. We test whether this corresponds to increased population growth rate in a range of glyphosate doses compared to the susceptible control strain throughout the course of adaptation to assess the level of resistance. Furthermore we test the hypothesis that there is an intrinsic cost to glyphosate resistance through continuous assaying of the population growth rate in the ancestral environment to assess whether the evolution of resistance is associated with a trade-off in performance. The expected possible outcomes here correspond to those set out in Fig 1.

## Methods

### Experimental design

#### Algal strain and chemostat setup

*Chlamydomonas reinhardtii* strain CC-1690 was obtained from the Chlamydomonas Resource Centre core collection. This strain has been previously used in experimental evolution studies, including for herbicide resistance evolution[28, 31, 32]. Prior to the experiment, the algae had been kept in static stock cultures for two years (Ebert algal medium, 25°C, 24h light), with batches being transferred to fresh medium fortnightly, then in continuous flow culture for two months under the experiment control conditions detailed below. All *C. reinhardtii* populations were then cultured in continuous flow through chemostats (see [39] for detailed protocol) in Ebert algal medium at a dilution rate of 0.15/day with a shared multichannel peristaltic pump (Watson-Marlow 205S/CA16) controlling the flow for all culture chambers, maintaining a culture volume of 380 ml *±*5%. The populations were kept on a light box providing white light from below and surrounded on all sides with white light fluorescent bulbs at a light level of 75 μmol m^-2^ s^-1^ 24 h a day. The cultures were heated by the light box to an internal temperature of 30°C, and kept in a controlled temperature room with an ambient temperature of 25°C. The cultures were continuously mixed by bubbling with air supplied from the building’s air supply taps. The 16 experimental *C. reinhardtii* populations were allowed 14 days after inoculation to reach steady state and acclimate to control conditions to ensure equal and high starting population sizes of approximately 2.5 *×* 10^5^/ml.

#### Herbicide concentrations and replication

10 out of the 16 populations were exposed to glyphosate (PESTANAL®, analytical standard) at a concentration of either 100 mg/L (lethal dose) and 50 mg/L (sublethal dose) 14 days after initial inoculation, with five chambers receiving each dose. 100 mg/L glyphosate completely inhibits population growth in batch cultures over at least 4 days[31, 32], whereas 50 mg/L glyphosate results in a population growth rate reduction. The remaining 6 chambers continued to receive the control medium.

### Population density and growth rate assays

The culture chambers were sampled daily from the middle of the culture to monitor population density through flow cytometry (Beckman Coulter CytoFLEX) starting 5 days after initial inoculation.

Growth assays were performed 1, 8, 22, 29, 36 and 43 days after glyphosate introduction. As the resistance trait is likely to be in the form of increased ability to grow in a given dose rather than complete insensitivity to glyphosate, i.e. a MIC will still exist, the resistance level was tested at five doses of glyphosate, ranging from sublethal to far above lethal for a naive strain and glyphosate resistance was defined as increased population growth rate in the presence of glyphosate compared to control population growth under the same conditions. Intrinsic trade-offs were assessed through comparing the population growth rate in the ancestral environment to the control population growth rate in the ancestral environment, defining a cost as a reduction in population growth rate under control conditions associated with increased resistance. A washed subsample from each population consisting of approximately 2.5 *×* 10^7^ cells was allowed to acclimate to the ancestral environment in 30 ml of control medium for 72 hours to allow at least one generation of growth without the herbicide treatment for cells permanently growth-inhibited by glyphosate to die and thus be removed from the subsample, as well as acclimation to the batch environment of the assays. The resistance and trade-off assays were carried out simultaneously with 10 μl of each washed sample added to 250 μl control medium containing 150, 125, 100, 75, 50 and 0 mg glyphosate/L respectively with 3 replicates of each. The fluorescence intensity (excitation: 485 nm, emission: 670 nm) for each sample was measured as a proxy for population density at 0, 6, 12 and 24 hours using a Tecan Spark 10M Multimode Microplate Reader. These time points adequately captured the log phase of growth both in control medium and under acute glyphosate stress in pilot studies. Between measurements, the assay populations were kept under the same temperature and light conditions as the chemostat populations.

### Population density through time analysis

All data were analysed using R (version 4.0.5) and CytExpert (Beckman Coulter). CytExpert was used to gate and count events detected in the PerCP-A channel (excitation: 488 nm, emission: 690/50 BP) to determine population density. This channel is used to detect chlorophyll *a* and represents a robust method for estimating algal density[40] which was further validated against manual haemocytometer counts for this system. Data from one of the control chambers and one of the 50 mg/L glyphosate chambers were removed after a malfunction (resulting in *n*=5 for controls and *n*=4 for 50 mg/L for the latter half of the data set). Growth-inhibition due to glyphosate was defined as a continuous decline in population density for at least 3 days following the addition of the herbicide. A continuous increase in population following this decline to eventually reach a steady state was considered evidence of a newly resistant population.

Day-to-day growth rate for each chamber was calculated as *GR* = *log*(*N*_*t*_ *− N*_*t*_ *−* 1)*/*Δ*t*, where *GR* denotes growth rate, *N* is the population density at a given time-point and the preceding time-point, and Δ*t* is the time elapsed between measurements. To test whether the day-to-day growth rate exhibited a pattern different from a flat fit, i.e. if there was a noticeable population decline and recovery coinciding with glyphosate treatment, R package `mgcv`[41] was used to run a hierarchical generalised additive model (hGAM) following[42]. The hGAM framework was chosen to allow modelling of the average treatment effect on the full time series while taking into account the differences between individual chamber populations, capturing the non-linear dynamics of the system. Day-to-day growth rate was modelled as a function of the interactions between time and glyphosate treatment as well as time and population. Populations within the same glyphosate treatment were assumed to have a similar functional form over time but with some intergroup variation and were thus fit with a shared smoothing parameter, whereas each glyphosate treatment was fit with independent smoothing parameters to allow for differences in wiggliness[42]. In both cases thin plate regression splines were used. Day was also included as a random effect smooth (equivalent to a random intercept) to account for day-to-day variation. The first 7 days of the experiment were excluded from the model to improve the fit as the population establishment phase is inherently different in its dynamics to the steady state and thus introduced unnecessary noise. The model residuals were also assessed for temporal autocorrelation within each chamber and were judged to be independent, and as such no correlation term was included in the model.

### Resistance level and fitness trade-off analysis

Times series were constructed of maximum growth rates for all replicates in the growth assays, where the growth rate for each day was the maximum growth rate measured within each 24h period of sampling. Mixed-effects models with this daily maximum growth rate as the response were fit with R `lme4` package[43] to assess the change in slope (i.e. resistance level) and intercept (i.e. associated cost in ancestral environment) throughout the course of resistance evolution for each population. Dose, day and treatment were fit as fixed effects and chamber as a random effect with varying intercept. Both growth rate and dose were scaled and centred to improve fit. R `emmeans` package[44] was used to estimate the dose response trends and intercepts obtained from the model.

As the populations never exhibited growth in 125 or 150 mg/L glyphosate and would reach a fluorescence intensity consistent with a population consisting of dead cells within 12 hours, these assays were excluded from the linear mixed-effects models to improve fit. The growth rate in these assays 43 days after glyphosate introduction were instead analysed as death rate assays using an ANOVA to determine if there was evidence for a higher growth rate (i.e. slower rate of death) to higher doses, suggesting increased tolerance associated with glyphosate resistance.

## Results

### Population density shows evidence of resistance 19 days into lethal dose treatment

While each population represents a separate evolutionary trajectory, there are clear overall patterns shared by the populations receiving the same treatment (Fig 2). All the populations receiving the lethal treatment exhibit a clear density decline where the day-to-day growth rate consistently averages below 0 for at least 3 consecutive days starting after the treatment was introduced, but the exact onset of this decline and its slope varies by chamber. All chambers at some point during the mortality phase have a growth rate around or below -0.15, i.e. the expected population density change if all cell growth and division is arrested while the population is diluted by the continuous inflow of media. This suggests that the populations are not just being diluted as cell division is suspended, but that cells are dying at a faster rate than division can take place. The mortality phase is immediately followed by a recovery phase where the growth rate consistently averages above 0 for several consecutive days, suggesting the population has evolved resistance. The earliest instance of this is 19 days after the treatment was applied, and all populations have entered the recovery phase by 22 days. This pattern is not apparent in the controls or sublethal glyphosate populations, and is reflected in the hGAM of the day-to-day growth rate.

**Figure 2:**
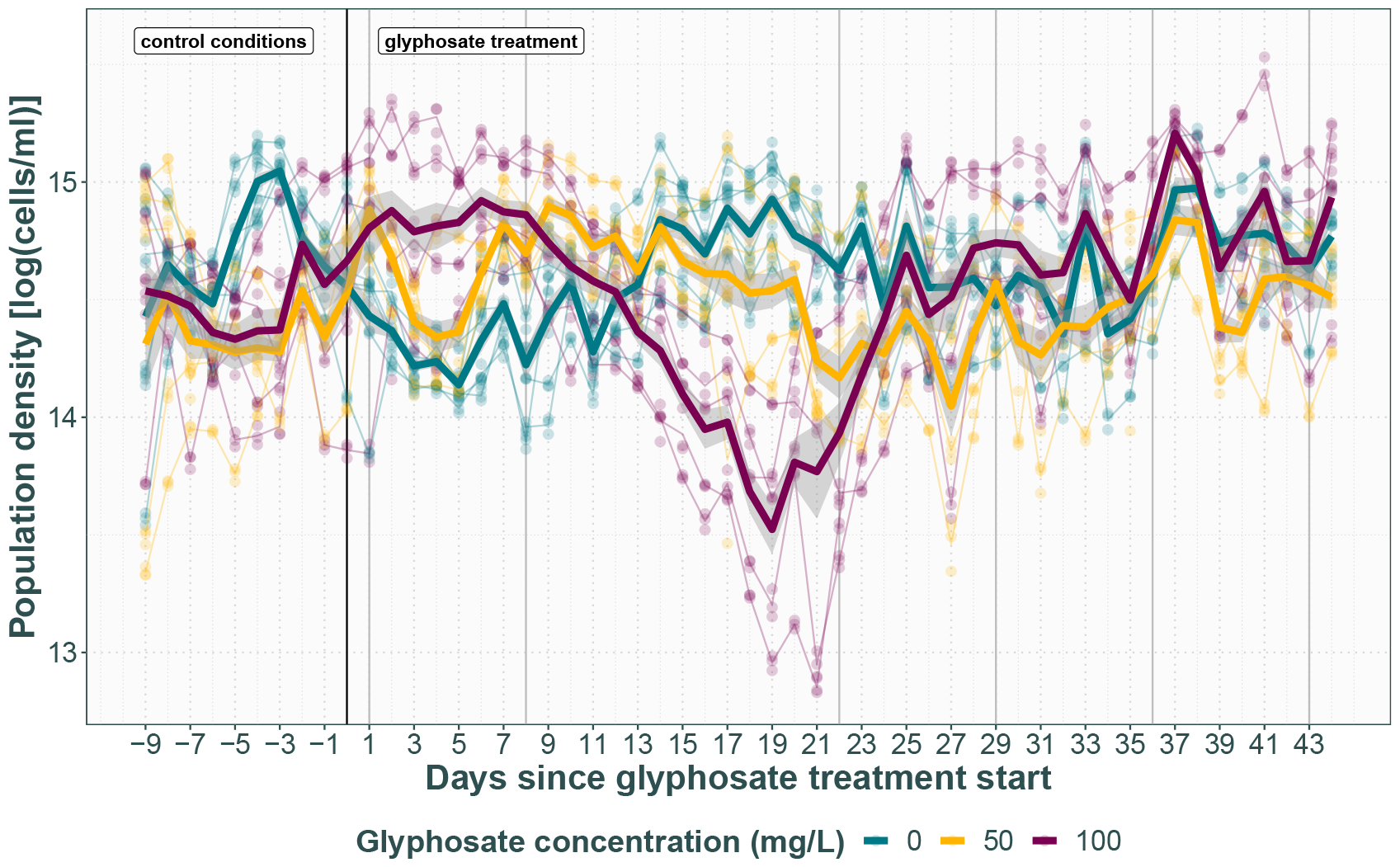
Population density throughout the experiment. Each point is a technical replicate, thin lines represent average chamber population density. Thick lines with SE represent average densities for the treatment groups. Grey vertical lines indicate growth assay sampling points.

The control populations and those receiving sublethal glyphosate treatment average day-to-day growth rates of zero after day 7, resulting in a flat fit by the hGAM (control F_8_=0, p=0.5; sublethal F_8_=0, p=0.7). By contrast, the model fit a smooth function significantly different from a flat fit for the populations receiving the lethal glyphosate treatment (F_8_=7.9, p<0.001).

### Growth assays show no evidence for increased level of resistance

43 days after the introduction of glyphosate there is no evidence of increased resistance to any of the tested doses or a shifted MIC, in direct contrast with the population density data indicating resistance has evolved 24 days prior. While there is an effect of dose (F_1_=454.1, p<0.001), treatment (F_2_=18.7, p<0.001) and day (F_5_=73.2, p<0.001), as well as their two-way (Dose*Treatment F_2_=21.1, p<0.001; Dose*Day F_5_=49.3, p<0.001; Treatment*Day F_10_=147.5, p<0.001) and three-way interactions (F_10_=21.7, p=0.02), on the growth rate, the dose-response relationships are indistinguishable between treatment lines Fig 3) by the end of the experiment Fig 4, Table 1). The intercepts of the dose response are also not significantly different between treatments, indicating no evidence of a fitness cost in the ancestral environment.

**Table 1:** Within-day pairwise contrasts of linear mixed effects model fixed effect term slopes and intercepts.

| Contrast |  | Slope |  |  | Intercept |  |  |
| --- | --- | --- | --- | --- | --- | --- | --- |
| Day | Treatments | df | t-ratio | p-value | df | t-ratio | p-value |
| 1 | 0 - 50 | 279.1 | 0.8 | >0.9 | 158.4 | -0.6 | >0.9 |
|  | 0 - 100 | 279.1 | 0.9 | >0.9 | 158.4 | -0.6 | >0.9 |
|  | 50 - 100 | 279.1 | 0.08 | >0.9 | 158.4 | 0.06 | >0.9 |
| 8 | 0 - 50 | 279.1 | -2.3 | 0.7 | 158.4 | 2.7 | 0.4 |
|  | 0 - 100 | 279.1 | -3.3 | 0.09 | 158.4 | 6.7 | <0.001 |
|  | 50 - 100 | 279.1 | -0.9 | >0.9 | 158.4 | 3.7 | 0.03 |
| 22 | 0 - 50 | 279.1 | -4.5 | 0.001 | 157.5 | 7.5 | <0.001 |
|  | 0 - 100 | 279.1 | -1.9 | 0.9 | 157.5 | 2.9 | 0.3 |
|  | 50 - 100 | 279.1 | 2.4 | 0.6 | 157.5 | -4.4 | 0.002 |
| 29 | 0 - 50 | 279.1 | -2.1 | 0.8 | 185.0 | 4.1 | 0.007 |
|  | 0 - 100 | 279.1 | -1.3 | >0.9 | 169.3 | 2.4 | 0.6 |
|  | 50 - 100 | 279.1 | 0.9 | >0.9 | 173.5 | -1.8 | >0.9 |
| 36 | 0 - 50 | 279.1 | -1.0 | >0.9 | 185.0 | 3.1 | 0.2 |
|  | 0 - 100 | 279.1 | 0.1 | >0.9 | 169.3 | -0.9 | >0.9 |
|  | 50 - 100 | 279.1 | 1.1 | >0.9 | 173.5 | -3.9 | 0.01 |
| 43 | 0 - 50 | 279.1 | -1.9 | 0.9 | 185.0 | 2.7 | 0.4 |
|  | 0 - 100 | 279.1 | -0.9 | >0.9 | 169.3 | -1.3 | >0.9 |
|  | 50 - 100 | 279.1 | 1.0 | >0.9 | 173.5 | -3.9 | 0.02 |

**Figure 3:**
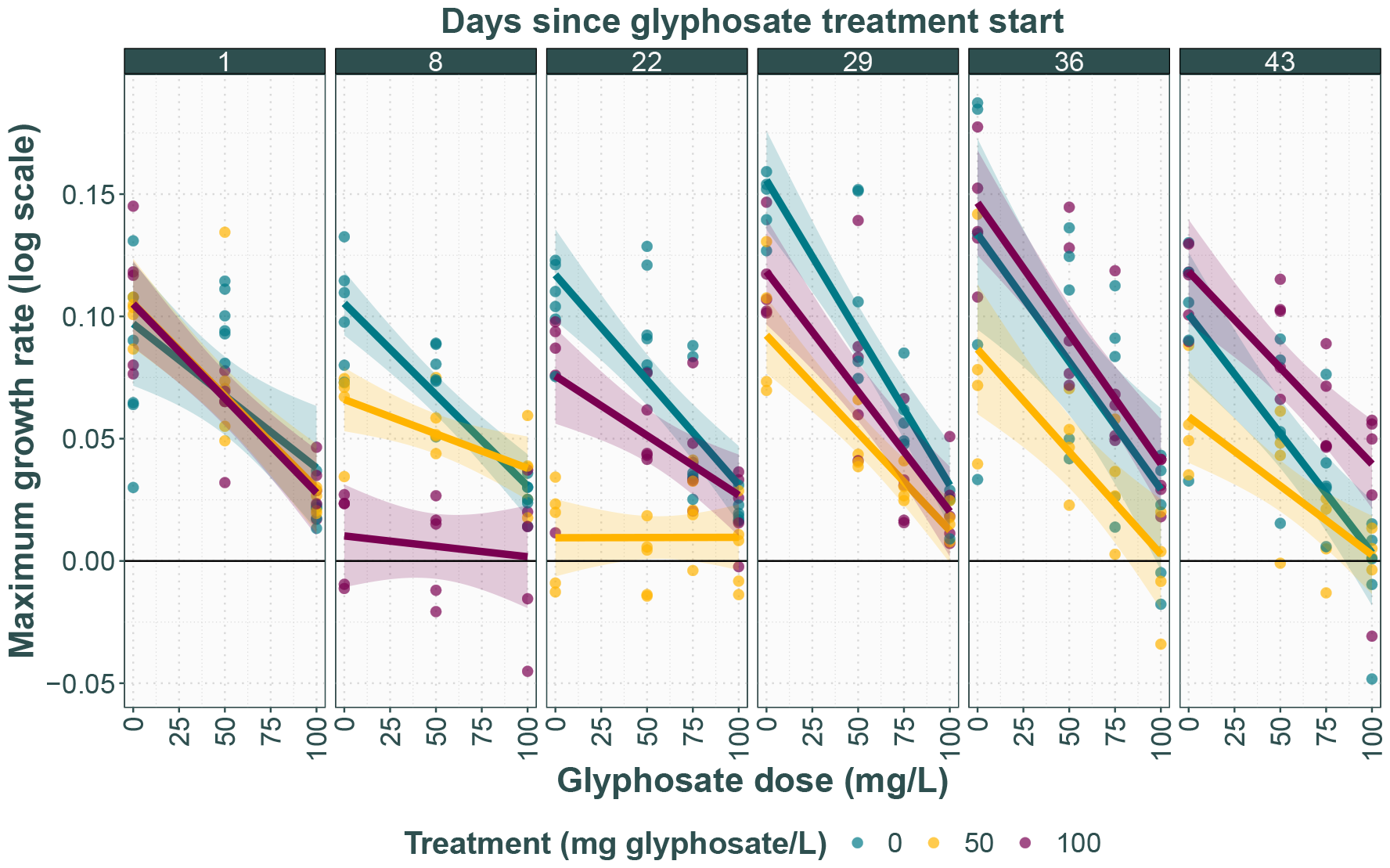
Maximum log growth rate in control medium and range of glyphosate doses for each growth assay. Each point is the average growth rate of a chamber, with lines and 95% confidence intervals representing the average for each treatment group.

**Figure 4:**
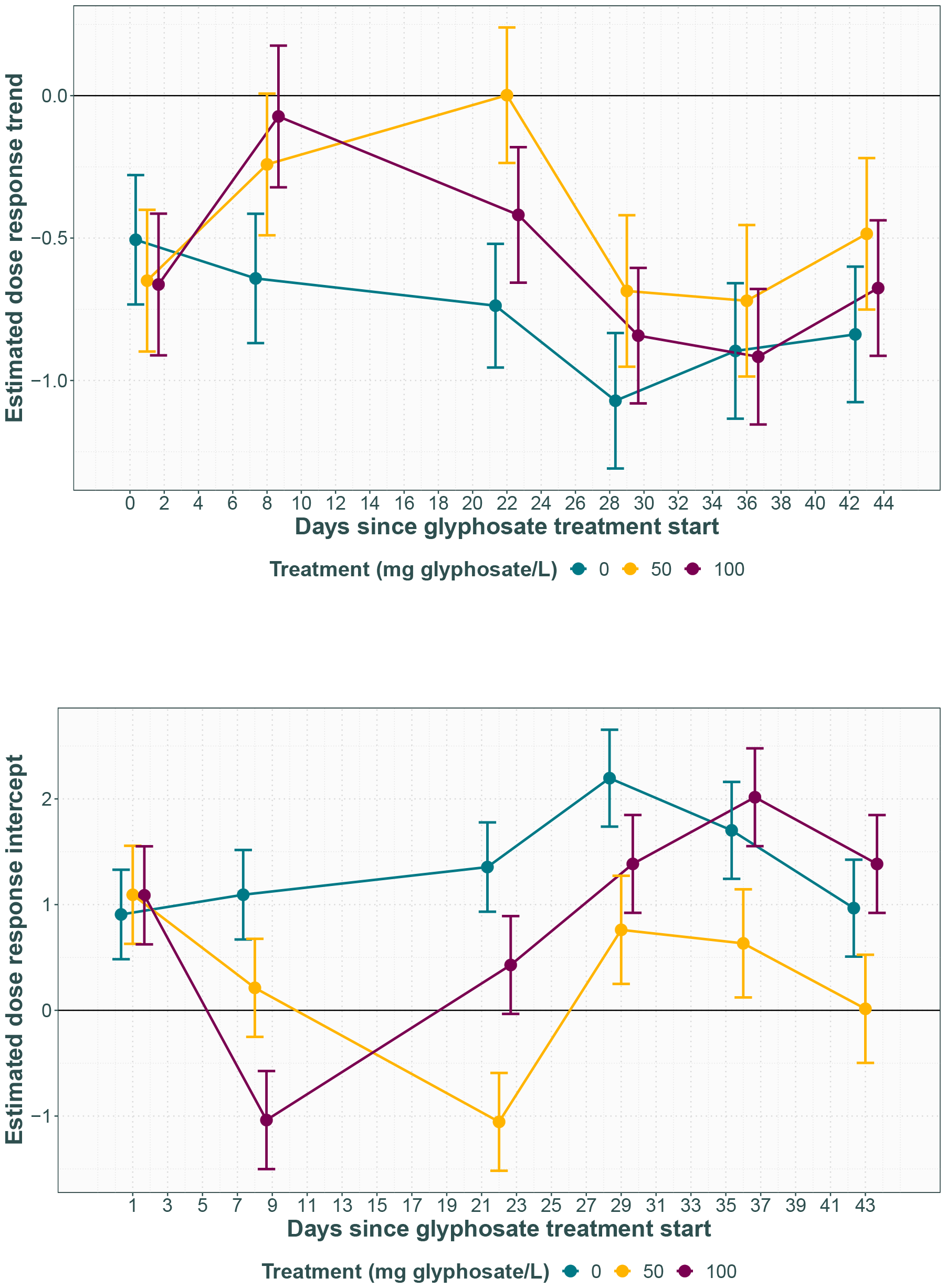
Dose response trend (top) and intercepts (bottom) with 95% CI through time as estimated by linear mixed effects model.

The death assays also reveal no differences in population performance between treatments at very high doses of glyphosate, with no effect of dose (F_1_=1.8, p=0.2), treatment (F_2_=2.5, p=0.1) or their interaction (F_2_=0.2, p=0.8) on growth rate. This further suggests no increased tolerance or shifted MIC as a result of either glyphosate treatment.

### Glyphosate treated populations show decreased initial assay performance in all doses

After comparable performance for all treatments right after the introduction of glyphosate, all glyphosate treated populations show a decrease in overall growth rate compared to the controls in the growth assays performed at 8 and 22 days post-glyphosate introduction Fig 4, Table 1. Concurrent with the population decline apparent in the population density for the lethal treatment, the slopes flatten and the intercepts decrease, reflecting decreased performance in all doses of glyphosate tested. This is followed by the slopes and intercepts returning to the same levels as those of the controls. Notably, while no evidence was found for a population density decline in the sublethal treatment populations, they exhibit the same slope and intercept patterns as the lethal populations, albeit lagging by roughly two weeks.

## Discussion

To manage resistance to economically important herbicides like glyphosate, we need to understand the underlying evolutionary dynamics. Model organisms such as green alga *C. reinhardtii* provide the opportunity to test the evolutionary theory under controlled lab conditions with huge population sizes and fast generation times, giving insight both to weed science and evolutionary biology[30, 45]. We here for the first time use chemostat cultures to evolve glyphosate resistance in *C. reinhardtii* to both a lethal and sublethal dose and monitor the effects on population density, resistance level and growth in the ancestral environment throughout the course of resistance evolution.

### Evidence for rapid resistance evolution

The population density data shows rapid evolution of resistance to glyphosate in populations experiencing a lethal dose, with the inflection point between the mortality phase and the recovery phase reflecting the point at which resistance becomes dominant appearing in the population as early as 19 days after the treatment was initiated. This is somewhat faster than the average pace of adaptation of 3.5 weeks for this system found in other studies [31, 33], likely owing to the larger population sizes and lack of evolutionary bottlenecks. The exact number of generations covered by this time is not possible to determine, as the growth inhibition of the majority of the population means growth rate is no longer determined by dilution rate as is otherwise expected in chemostat systems [46]. However, given the light intensity used, the generation time is likely between 24–48h [47], suggesting resistance evolved in less than 20 generations.

By contrast, the dose response relationship by the end of the experiment for this treatment line does not indicate a shift in the MIC as it is not significantly different from the control treatment. Instead we see a marked decrease in growth rate at all glyphosate doses as well as in the absence of glyphosate coinciding with the morbidity phase, followed by a recovery. This is likely due to the populations comprising multiple, co-existing genetic lineages with different adaptive strategies, a consistent finding in evolution experiments using chemostats[46, 48]. This population structure is largely attributed to soft selective sweeps and proportion fluctuations being more common than hard sweeps to fixation of any allele[46]. As such, when measuring overall population performance, as opposed to isolating and testing labelled clones, the result is an average of those genetic lineages, and the performance of individual lineages is obscured. A new, fitter genotype will only have a noticeable signal in the assays once it has reached a larger proportion of the population. Thus, there is likely to be a lag between the emergence of a new lineage and its detectability in the growth assays as performed here, and the coexistence of less fit lineages means the true fitness of any new mutation is likely to be underestimated. Furthermore, cells that are growth-inhibited by the glyphosate will remain in the population until they die or are flushed out, and these cells will lower the average population performance in the assays. A considerable proportion of the population being growth-inhibited at the beginning of the experiment may be why the treated lineages perform poorly in the growth assays, while growth is indeed detectable in the population density measurements.

The sublethal glyphosate dose treatment appears to have a negligible effect on population density, likely due to the dilution rate allowing a moderate reduction in growth rate without resulting in over-dilution or the model lacking the statistical power to pick up subtler effects. While there is no evidence for a shifted MIC in the growth assays, the decrease in performance with a subsequent recovery suggests the populations have evolved. This pattern lags behind that of the lethal dose by roughly two weeks, consistent with the sublethal dose constituting a weaker selective pressure providing an alternate path to resistance through accumulation of minor gene traits to result in moderate resistance[12, 13].

### Implications for intrinsic costs and the resistance mechanism

For a reduction in growth rate in the ancestral environment to represent the signal of a likely trade-off with glyphosate resistance, it should be observed in conjunction with increased resistance. While the evolution of increased resistance to the lethal dose treatment as evidenced by the population density data must necessarily reflect a shift of the MIC, the fact that the dose response trends include a strong signal of the growth-inhibited portion of the population makes it difficult to draw conclusions about any associated fitness costs. However, the fact that the estimated intercept for the lethal treatment populations is not significantly different from the control populations by the last assay suggests that the fitness cost of resistance in the ancestral environment is either small or non-existent. This is also the case for the sublethal dose populations as their evolution of resistance is only inferred by exhibiting a similar pattern in the growth assays. As two of the previous studies examining the fitness costs of glyphosate resistance in *C. reinhardtii* found evidence of a minor[31] and major[33] growth rate reduction in the ancestral environment associated with evolved resistance, and the third found a positive correlation between resistance level and fitness[28], our results suggest that fitness costs are not universal in this system and that the resistance trait in all studies to date may be conferred by different molecular mechanisms.

Given the growth-inhibiting action of glyphosate, the resistance mechanism for the lethal dose populations is likely initially based on standing genetic variation rather than *de novo*-mutations, which has also been found to be important for glyphosate resistance evolution in some weeds[49, 50, 51]. While mutations may have accumulated subsequently to improve resistance, the short time frame of the experiment limits the likelihood of polygenic resistance mechanisms. Furthermore, sequential point mutation genotypes changing the structure of EPSPS tend to render it highly insensitive to glyphosate and confer very high resistance[52, 53, 54, 55], meaning we would then have expected to see a reduction in death rate, if not evidence of growth, at the very high doses of glyphosate tested in the death assays. A single substitution causing a structural change to EPSPS may instead cause a moderate resistance increase at variable cost depending on species[56, 57, 58, 59, 60]. Incidences of these genotypes are expected to be rarely occurring as part of standing genetic variation as the structure of EPSPS is highly conserved, although they may arise quickly under strong directional selective pressure[1]. An amplification of EPSPS[61, 62], however, may possibly be present already in a large population [1], while conferring moderate resistance[63] to no detectable resistance cost[27]. Similarly, resistance mechanisms affecting other parts of the cellular machinery to reduce the dose reaching EPSPS such as increased vacuolar sequestration[25, 64, 65, 26] or glyphosate degradation[66] could theoretically have evolved in only a few generations, resulting in a medium level of resistance without obvious intrinsic fitness consequences[1].

While the lethal and sublethal dose treatments appear to be following the same growth assay pattern, the respective treatments are likely selecting for different fitness optima and may thus be selecting for different resistance mechanisms. Most cases of herbicide resistant weeds documented in the field exhibit major single gene mechanisms of resistance, due to high herbicide doses only selecting for sufficiently high resistance mutations[1]. Lower doses instead select for a range of mutations conferring a greater variety in level of resistance, and previous studies indicate that sublethal doses may favour accumulation of minor resistance traits as well as polygenic traits[12, 13, 67]. However, it has also been suggested that the chronic stress of low herbicide doses might lead to increased mutation rates, and thus increase generation of large impact mutations[11, 14, 68].

### Broader implications and future research

While this experiment covers only the initial dynamics of glyphosate resistance evolution in *C. reinhardtii*, these early stages are likely to determine the later outcome of the process[5] and the application of chemostats to the chosen system allowed tight control of the specific selective pressures[46] as well as monitoring the dynamics through time. These results also add to the increasing number of studies emphasising the potential *C. reinhardtii*, along with other algae, have for a rapid evolutionary response to herbicides, demonstrating their usefulness as a link between weed science and evolutionary biology [31, 32, 28, 33].

The results reported here also have direct implications for wild *C. reinhardtii* and other algae as agricultural runoff has led to glyphosate being a widely occurring pollutant of non-target ecosystems[16]. As the dose reaching non-target ecosystems will vary, the indication that a sublethal dose at half the MIC may still lead to relatively rapid resistance evolution is of particular interest. Further investigation of the effects of even lower doses could determine how dose affects both the pace of evolution and the particular resistance trait selected for, both which may determine the overall outcome for the ecosystem[36, 37].

Future research should also investigate the evolutionary dynamics of longer term effects of glyphosate exposure as further adaptation and refinement of the resistance mechanism is likely. Increased replication would also allow for refining the hGAM to add in more terms to explain variation, e.g. to control for perturbations to the steady state caused by sampling events, thus creating a more sensitive model able to detect subtler effects of lower doses of glyphosate. Furthermore, isolation of the genotypes present in the evolving populations along with competition assays would allow detailed characterisation of the resistance level and associated fitness costs, whereas -omics analyses could reveal whether sublethal and lethal doses select for different resistance mechanisms or merely provide different paths to the same goal.

