## Supplemental material 1 for "Glyphosate resistance evolution to lethal and sublethal doses in chemostat populations of model organism *Chlamydomonas reinhardtii*"

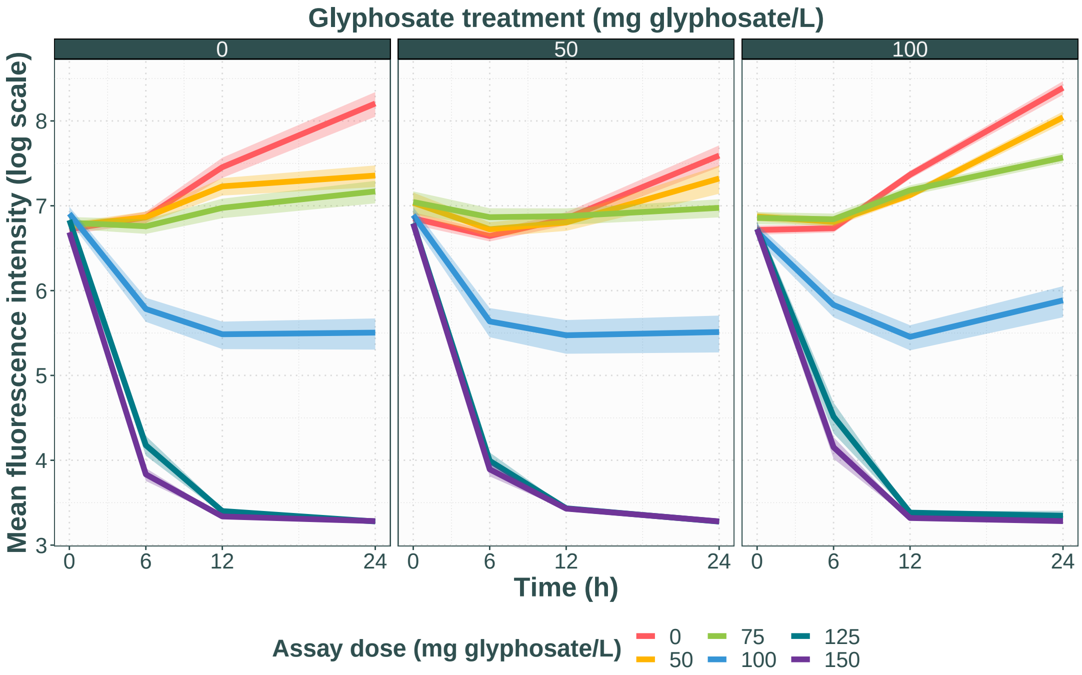


Figure 1: Mean fluorescence intensity, a proxy for population density, through time in the growth assay 43 days after glyphosate introduction. Growth in each dose is represented as a separate colour. Each selection treatment is represented as a separate panel (0, 50, 100 mg/L glyphosate).

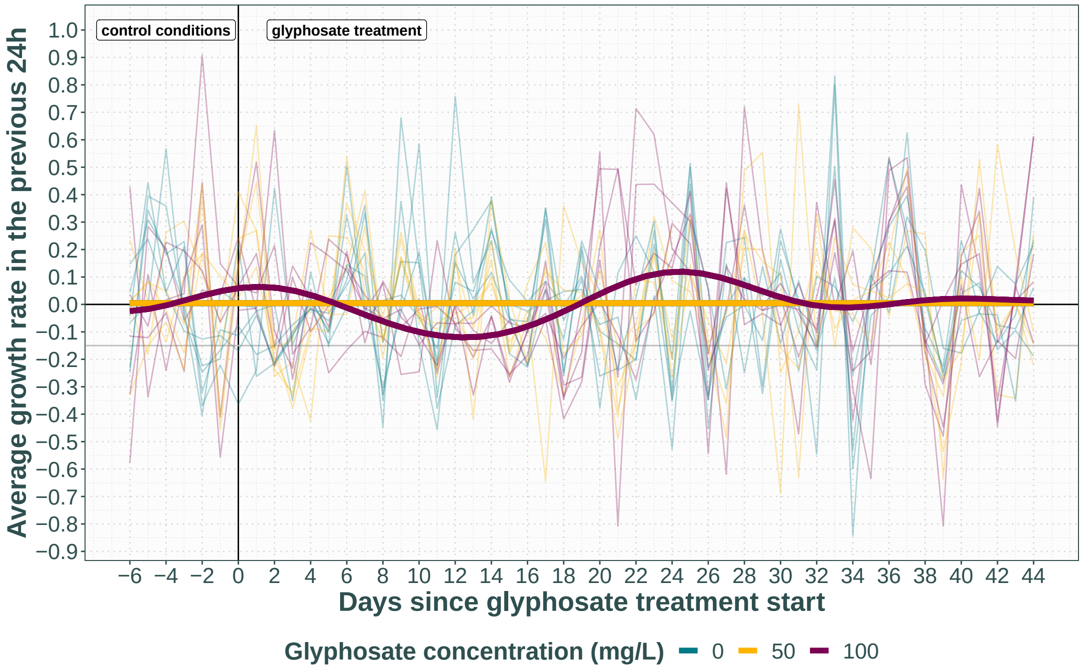


Figure 2: Growth rate between days 8 and 57 of the experiment. Each thin line represents a population, thick lines display the predicted fit for each treatment by the hGAM. The horizontal grey line represents the dilution rate of 0.15.

Table 1: Output for hGAM of day-to-day growth rate. edf refers to the effective degrees of freedom, i.e. the complexity of the smooth. The Ref.df, F and p-value are generated by an ANOVA to test the overall significance of the smooth, i.e. testing if it is different from a flat fit.

| **Smooth term** | **edf** | **Ref.df** | **F** | **p-value** |
|---|---|---|---|---|
| s(exp_day):treatment_group0 | 0.000002 | 8 | 0 | 0.5 |
| s(exp_day):treatment_group50 | 0.000009 | 8 | 0 | 0.7 |
| s(exp_day):treatment_group100 | 5.7 | 8 | 7.9 | 0.001 |
| s(treatment_group) | 0.0000001 | 2 | 0 | $< 0.9$ |
| s(exp_day,chamber) | 0.00001 | 158 | 0 | $< 0.9$ |
| s(dayID) | 41.5 | 50 | 5 | $< 0.001$ |

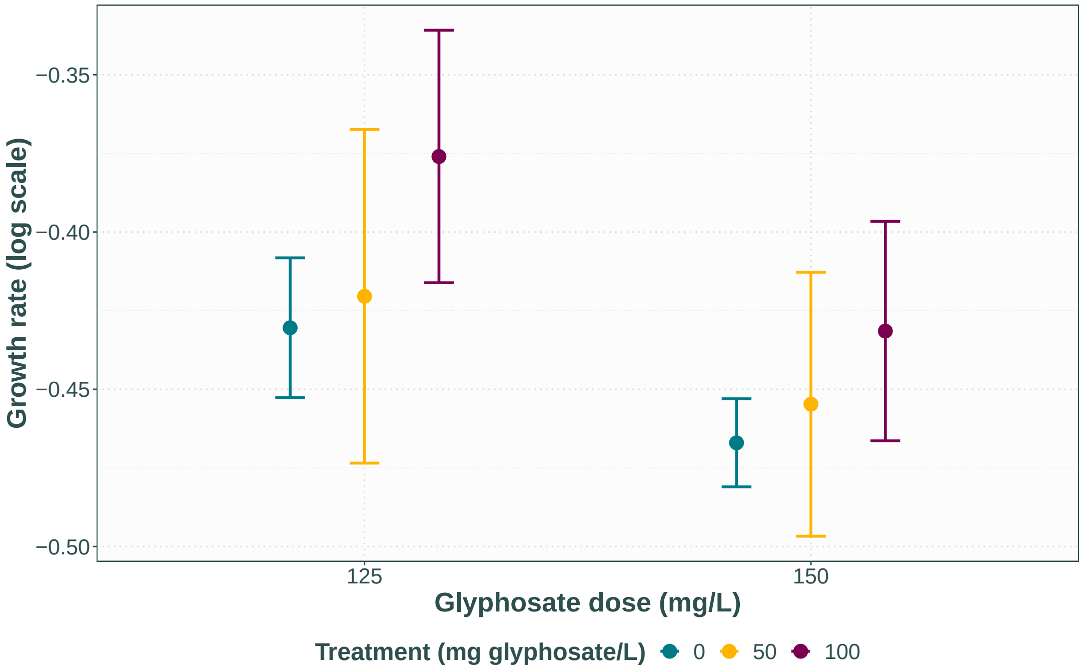


Figure 3: Average growth rate over 6 hours in very high lethal doses of glyphosate 43 days after glyphosate introduction.
